# Replication fork remodeling proteins, Smc5/6 and Rtt107, promote palindrome-mediated genome instability

**DOI:** 10.64898/2026.04.29.721645

**Authors:** Alex B. Costa, Anissia Ait Saada, Thy Pham, Jiayi Fan, Rebecca Lever, Xiaolan Zhao, Kirill Lobachev

**Affiliations:** Georgia Institute of Technology, School of Biological Sciences, Atlanta, Georgia, USA; Memorial Sloan-Kettering Cancer Center, Molecular Biology Program, New York, New York, USA

**Keywords:** Palindromes, secondary-structures, cruciform, hairpin, double-strand breaks, structure-specific nucleases, chromosome fragility

## Abstract

Palindromic sequences are a potent source of genomic instability that can lead to cancer and hereditary diseases in humans. Molecular evidence shows that palindrome instability results from the formation of secondary structures, such as hairpins and cruciforms, which are cleaved by structure-specific nucleases. However, the mechanisms underlying cruciform formation and cleavage at palindromic sequences in eukaryotic cells remain incompletely understood. Here, we describe a pathway for secondary structure formation and chromosomal breakage at palindromes involving DNA helicase Mph1, Rad51 recombinase, Rad54 ATPase DNA strand remodeler, Rtt107 scaffold protein, and the multifunctional Smc5/6 complex. Deletion or mutation of any of these components results in a similar reduction in double-strand breaks at an *Alu* palindrome and a significant decrease in chromosomal rearrangements. We propose that Mph1, Rad51, and Rad54 work together at stalled replication forks to generate cruciform structures via fork remodeling, while Smc5/6 and Rtt107 mark the cruciforms, indicating an appropriate substrate for nuclease cleavage. As members of this pathway are conserved in humans, the uncovered mechanisms may underlie genomic instability at palindrome sites involved in disease etiology.

## Introduction

Palindromes are self-complementary DNA sequences that are symmetrical around a central point and, through intrastrand base pairing, can form hairpin and cruciform structures. These structures can block replication and are targeted by multiple structure-specific nucleases, resulting in a double-strand break (DSB). Aberrant repair of DSBs can cause gross chromosomal rearrangements (GCRs), including large deletions, insertions, duplications, inversions, and translocations, which may lead to the loss, gain, or dysregulation of genetic information. Palindromic DNA sequences are found throughout the human genome (1) and are known as hotspots for chromosomal breakage and rearrangements that drive tumorigenesis and multiple hereditary diseases (2–13). Palindromes may also play roles in other diseases, as disease-associated risk variants are 14 times more likely to be located at palindromic sequences than elsewhere in the genome (1). In all cases, pathogenic palindrome-mediated chromosomal rearrangements are initiated by secondary structure formation and nucleolytic cleavage. Therefore, identifying pathways of secondary structure formation and their recognition and resolution is essential for understanding disease etiology.

Perfect palindromes and even inverted repeats separated by spacer regions can efficiently form hairpins within single-stranded DNA (ssDNA), such as during discontinuous synthesis on the lagging strand of the replication fork, during break-induced replication, or in resected regions caused by telomere uncapping (14–16). Hairpin formation during DNA replication can cause fork arrest and trigger nuclease attack. In E. coli, the SbcC/D nuclease cleaves hairpins that halt replication fork progression (17). Previously, we showed that in budding yeast, the Mre11/Rad50/Xrs2 (MRX) nuclease, the eukaryotic homolog of SbcCD, also cleaves hairpins, allowing invasion of the nascent sister chromatid to initiate repair. However, hairpins evade MRX-mediated cleavage, as we still observe strong replication arrest in strains where MRX is active (18). Although not yet demonstrated for hairpin-mediated blockage, replication fork regression and template switching are two processes that may help overcome lesions blocking replication. First, during fork reversal, the two nascent DNA strands anneal, enabling the fork to regress into a four-way junction, known as a “chicken foot” structure (19,20). This stabilization allows for replication restart without chromosomal breakage. Second, template switching is a process where a nascent DNA strand blocked by a lesion invades the nondamaged sister chromatid, using it as a template to bypass the lesion (21).

Cruciform formation in double-stranded DNA requires an opening of 8-10 bp at the center of symmetry and costs energy, which can be supplied by negative supercoiling. In *E. coli,* cruciforms readily form in negatively supercoiled plasmids (22). The exact mechanisms that drive cruciform extrusion in linear eukaryotic chromosomes are not yet known. *In vitro* studies suggest that negative DNA supercoiling may be generated by chromatin remodeling complexes (23) and DNA repair ATPases (24,25), promoting the formation of cruciform structures from linear DNA molecules. It has also been proposed that negative superhelicity generated during transcription can contribute to cruciform extrusion (26,27). In budding yeast, using a conditional-palindrome assay, we showed that two-ended hairpin-capped symmetrical breaks, indicative of cruciform formation, and resolution accumulate during the cell cycle in cells going through S-phase and progressing to G2. We also observed that symmetrical breaks occur in nocodazole-arrested nondividing cells (18). These findings suggest that cruciforms can form via two pathways: one requiring DNA synthesis and another that is replication-independent and operates in G2 cells.

Here, an unbiased genome-wide mutagenesis screen in *S. cerevisiae* was performed to identify new factors involved in palindrome-induced chromosomal instability. We observed that mutations in the DNA helicase Mph1, the Rad51 recombinase, the Rad54 repair ATPase, the Rtt107 scaffold protein, and Smc5/6 reduce DSB formation at an *Alu* palindrome. Our two-dimensional gel electrophoresis analysis of replication intermediates indicates that the recombination machinery promotes cruciform formation at the replication fork through template switching and an attempt to bypass hairpin-mediated blockage. We demonstrate that although the Smc5/6 complex does not participate in template switching or cruciform formation, it acts within the same pathway of palindrome instability and, along with Rtt107, may mark cruciforms at the replication fork as targets for nuclease attack. This research uncovers a new pathway composed of highly conserved proteins for cruciform formation and processing in eukaryotic cells.

## Results

### An experimental system to monitor DSB formation at palindromic repeats and used to identify hypo-fragility mutants

The experimental system for monitoring GCR induction and DSB formation at the *Alu* palindrome (*Alu*-PAL) is based on the loss of the *CAN1* and *ADE2* genes and the amplification of the *CUP1* and *SFA1* genes relocated to chromosome VI (Supplemental Figure 1). This assay for detecting GCRs and gene amplification is similar to those developed for chromosome V and described in (18,28). Moving the *Alu*-PAL, GCR cassette (*CAN1-ADE2*), and the gene amplification cassette (*SFA1-CUP1*) to chromosome VI makes the gene amplification assay more specific to the DSB outcome. We found that, in addition to gene amplification resulting from *Alu*-PAL-mediated breakage, non-disjunction of chromosome V can also contribute to the appearance of copper- and formaldehyde-resistant colonies (28). It has been shown that disomy of chromosome VI is lethal because of the presence of dose-sensitive *TUB2* and *ACT1* genes (29). Therefore, placing the *SFA1-CUP1* cassette on the right arm of chromosome VI enables the specific selection of extrachromosomal gene amplification events caused by *Alu*-PAL-mediated hairpin-capped breaks. The *LYS2* gene, containing *Alu*-PAL, was inserted at ChrVI::253403, 17 kb from the right telomere. *CAN1-ADE2* and *SFA1-CUP1-hphMX* cassettes were placed telomeric to *LYS2*, so the region between the *Alu*-PAL and the telomere, which lacks essential genes, extends approximately 29 kb. *Alu*-Pal-induced GCRs are selected as canavanine-resistant red (Ade^-^) colonies, while *Alu*-Pal-induced gene amplification events are manifested as copper-and formaldehyde-resistant colonies. The deduced mechanism for palindrome-induced GCRs is the formation of hairpin-capped breaks that result in dicentric dimers, followed by breakage later in cytokinesis, which requires repair, often involving break-induced replication using a non-homologous chromosome as a template. Gene amplification events are characterized by the accumulation of extrachromosomal acentric dimers, which result from hairpin-capped breaks, duplication of the broken molecule, and random segregation of centromere-lacking inverted dimers (28).

### Genome-wide screen identifies Mph1 as a promoter of palindrome-mediated instability

Previously, we found that Mre11/Rad50/Xrs2 (MRX) and Mus81/Mms4 endonucleases are involved in generating DSBs at palindromic sequences, where MRX and its nuclease regulator Sae2 cleave hairpins during replication, while Mus81/Mms4 targets cruciforms at transcribed repeats (18). However, hairpin-capped breaks are readily detectable at *Alu*-palindromes in *sae2Δmus81Δ* strains, indicating that other nuclease(s) can cleave cruciform structures in yeast. Additionally, we found that cruciforms accumulate during S-phase and are formed in G2 in non-dividing cells through unknown mechanisms (18). Therefore, using the system described above, we started a screen in *sae2Δsmus8Δ1* strain for mutants with reduced fragility, aiming to identify 1) a nuclease responsible for cruciform cleavage; 2) proteins that promote cruciform formation; and/or 3) regulators of nuclease or cruciform-forming activities.

Cells were treated with ethyl methane sulfonate (EMS) to induce genome-wide point mutations. Since these cells were DNA repair-deficient, a low EMS concentration (as described in Materials and Methods) was used to reduce mortality, the number of mutations, and to simplify downstream analysis. Approximately 113,000 colonies were screened for mutants with decreased arm loss and gene amplification levels. 128 mutants exhibited a hypo-GCR phenotype in both assays. Among these, two also showed reduced DSB formation. Next-generation sequencing (NGS) identified *mph1-292T>C* (*mph1-S98P*) as responsible for the hypo-fragility phenotype (an example of this analysis is presented in Supplemental Figure 1). This mutation is located in the ATP-binding domain of the Mph1 DNA helicase (Uniprot database). Additionally, *mph1-R604H* was identified as another hypo-fragile allele, residing in the helicase C-terminal domain (Uniprot database). Mph1 is an ATP-dependent, 3’-5’ helicase essential for maintaining genomic stability and is homologous to the human Fanconi Anemia protein FANCM (30).

### Mph1 is involved in both promoting palindrome-mediated breaks and the repair of broken chromosomes

To validate Mph1 as a factor in palindrome fragility and to confirm that the impact of *mph1* mutations does not depend on the context, we measured GCR rates and DSB formation in strains carrying the GCR cassette on chromosome V (18), Figure 1A). This assay on chromosome V is also convenient because replication timing and conditions for DSB detection have been previously determined and optimized (18).

**Figure 1.**
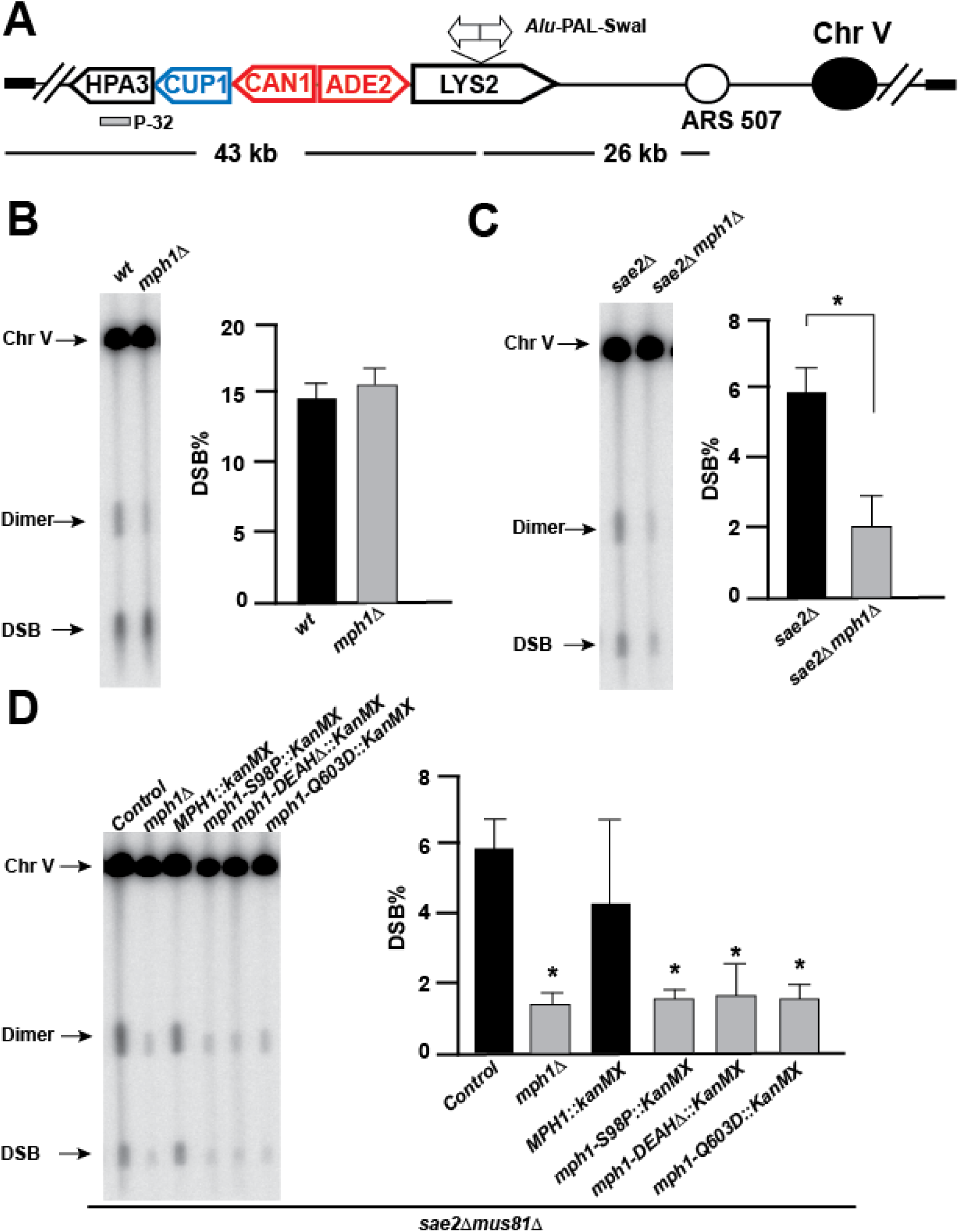
Mph1 helicase contributes to palindrome-mediated DSBs. A. Scheme of *Alu*-PAL and GCR cassette located on chromosome V. Secondary structures at *Alu*-PAL lead to the formation of hairpin-capped breaks (∼43 kb), and duplication of DSB fragment results in an inverted dimer twice the size of the break (∼89 kb). DSBs can be detected by Southern blot and hybridization after 26 hours of chromosome separation by CHEF. The *HPA3* probe (grey rectangle) is used to highlight DSBs, dimers, and intact chromosome V. B. C. and D. Representative images of palindrome-mediated DSB detection by Southern blot hybridization in wild-type, *sae2*Δ, and *sae2*Δ *mus81*Δ cells with *MPH1* and *mph1* mutants. Quantifications of DSBs are shown next to the gel images. Values represent the mean of at least 3 independent biological replicates ± standard deviation; * p < 0.05. P-values were calculated by comparing *mph1* mutants to their matched genetic controls. Control in D indicates *sae2Δmus81Δ* double mutant. Wild type *MPH1* and mutant *mph1* alleles were brought as a part of *kan*MX cassette.

The Mph1 was knocked out in wild-type (WT), *sae2*Δ, and *sae2*Δ*mus81*Δ backgrounds, and DSBs and GCRs were measured in these mutants. In WT cells, deleting *MPH1* did not reduce DSBs (Figure 1B). We previously showed that MRX/Sae2 plays a dual role in IR-induced DSB metabolism: it accounts for roughly 30-50% of breaks at perfect palindromes and is necessary for processing hairpin-capped breaks resulting from cruciform resolution (18). Therefore, the lack of a detectable impact of *mph1*Δ on DSB formation in WT is expected, because MRX/Sae2-mediated and processed breaks create a strong background of DNA intermediates that mask the effect. However, deleting *MPH1* resulted in about a 10-fold reduction in GCRs (Table 1). In *sae2*Δ and *sae2*Δ*mus81*Δ backgrounds, deletion of *MPH1* caused an approximately 3-fold decrease in DSBs, lowering the level of DSBs below 3% (Figure 1C, D). In *sae2*Δ or *sae2*Δ*mus81*Δ cells, deleting *MPH1* also led to roughly a 5- and 8-fold reduction in GCRs, respectively (Table 1). The greater impact of *mph1*Δ on GCRs than on DSBs suggests that it not only affects palindrome breakage but also plays a role in repairing breaks.

**Table 1:** Deletion of *MPH1* reduces palindrome-mediated GCRs.

| Strain | GCR rate x 10<sup>-7</sup> |
| --- | --- |
| Wild type | 6,109 (4,927-6,705)* |
| *mph1Δ* | 533 (458-595) |
| *sae2Δ* | 6,548 (5,029-8,030) |
| *sae2Δ mph1Δ* | 1,283 (1,013-1,582) |
| *sae2Δ mus81Δ* | 4,713 (4,142-6,382) |
| *sae2Δ mus81Δ mph1Δ* | 694 (658-841) |

\*Numbers in parentheses correspond to the 95% confidence interval

To determine which attributes of the Mph1 protein are required for palindrome instability, we tested three mutants. These included i) *mph1-S98P* uncovered from our screen, ii) *mph1-DEAH*Δ lacking the DEAH motif required for ATP-binding, and iii) mph1-Q603D that mutated a key residue for its helicase activity (32). We found that all three mutants mimicked the phenotype of *mph1*Δ, indicating that the helicase activity itself is crucial for Mph1-mediated effects on palindrome stability (Figure 1D).

### Mph1-interacting proteins Rad51, Rad54, Rtt107, and Smc5/6 are involved in promoting palindrome fragility

Mph1 is a 3’-5’ DNA helicase involved in various biological processes, including error-free bypass of DNA lesions, inter-strand cross-link repair, recombinational repair, and replication fork remodelling (33). To better understand Mph1’s role in palindrome stability, we examined Mph1 interacting partners that play a role in different DNA repair pathways. These non-essential genes were knocked out in the *sae2*Δ*mus81*Δ background, and DSBs were measured by Southern blotting. Deletion of *RAD51*, *RAD54*, and *RTT107* caused a significant reduction in DSBs, with Rad51 and Rad54 contributions nearly as important as Mph1 itself (Figure 2A).

**Figure 2.**
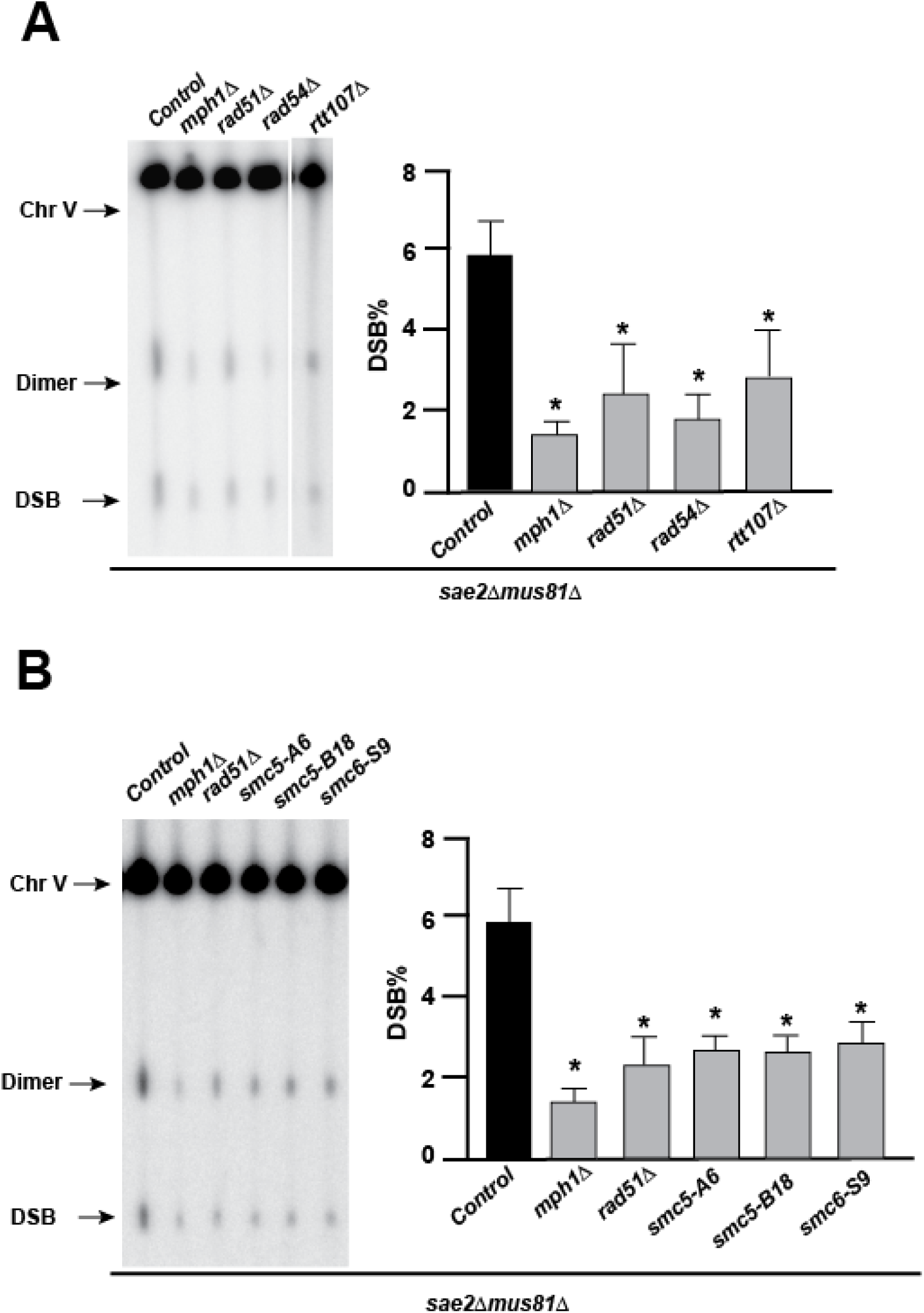
Rad51, Rad54, Rtt107, and Smc5/6 promote palindrome instability. A. Representative image of palindrome-mediated DSB detection by Southern blot hybridization of *MPH1*, *RAD51*, *RAD54* and *RTT017* knockout cells in a *sae2*Δ *mus81*Δ genetic background. Quantifications of DSBs are shown next to the gel images. B. Representative image of palindrome-mediated double strand break detection by Southern blot hybridization of cells without *MPH1* or *RAD51* or containing *smc5-A6*, *smc5-B18* and *smc6-S9* alleles in a *sae2*Δ *mus81*Δ genetic background. Quantifications of DSBs are shown next to the gel images. Values are the mean of at least 3 independent biological replicates ± standard deviation; * *p* < 0.05. *P-*values were calculated by comparing mutants to their matched genetic control.

Mph1 and Rtt107 interact with Smc5/6 both *in vitro* and *in vivo* (34–38). This prompted us to investigate whether mutations in the Smc5/6 complex have a similar effect on palindrome fragility as deficiencies in Mph1 and Rtt107. *SMC5::URA3* and *SMC6::URA3* fragments were amplified using mutagenic PCR, and the resulting libraries were used to transform the *sae2*Δ*mus81*Δ strain containing the *Alu*-PAL assay on Chr V. Three mutants—*smc5-A6*, *smc5-B18*, and *smc6-S9*—showed a 2-3 fold reduction in palindrome-mediated DSBs and had DSB levels similar to those observed with *MPH1* or *RAD51* deletion (Figure 2B).

### Conserved Smc5/6 features affect palindrome fragility

Sequencing of the above *smc5* and *smc6* alleles revealed that each contained multiple mutations distributed across the head (ATPase domain), arm, and hinge regions (Supplemental Figure 2A). Because these alleles contained multiple changes, we aimed to identify the most impactful change(s) within each allele. Guided by cryo-EM structures of the budding yeast Smc5/6 and sequence alignments of the proteins, we evaluated the potential effects of the mutations on local structural properties and residue conservation (Supplemental Figure 2A). Among residues changed in the *smc5-A6* allele, L897 shows high similarity among Smc5 homologs across species, and the L897Q change can alter the hydrophobicity of the surrounding region of the protein. For mutations within the s*mc5-B18* allele, we focused on *smc5-C20*Δ. The deleted region interacts with the Nse4 subunit (Supplemental Figure 2B), which could support the stability of the complex, although its sequence is not highly conserved (Supplemental Figure 2C). For the *smc6-S9* allele, we focused on P1038L, as P1038 is required to make a turn in the peptide chain located between the C-loop and Walker B motif of the ATPase domain (Supplemental Figure 2D). As such, its mutation to Leu has the potential to disrupt folding and influence the ATPase function. Although P1038 is not a highly conserved residue, Smc6 homologs in other species all contain a proline in proximity that could achieve the same structural effect (Supplemental Figure 2E).

We first assessed the three mutations for their effects on palindrome-mediated DSBs. While *smc5-L897Q* did not exhibit a defect, suggesting that more than one mutation is needed for the phenotype, *smc5-C20*Δ exhibited significantly reduced palindrome-mediated DSBs. Furthermore, smc6-P1038L was found to be both necessary and sufficient to reproduce the effect observed for *smc6-S9*, leading to approximately a 2.5-fold reduction in DSBs (Figure 3A). Given that *smc5-C20*Δ and *smc6-P1038L* exhibited reduced palindrome-mediated DSB levels, we assessed whether they could also affect palindrome-mediated GCR levels. In the *sae2*Δ *mus81*Δ background, we found that either mutation led to a 2.5-fold decrease in GCRs (Figure 3B). These results show that both Smc5 and Smc6 promote fragility at palindromes, and that the C-terminal regions of each protein are important for this function. Since the s*mc5-C20*Δ mutation has the potential to affect Nse4 interaction (Supplemental Figure 2B) (39), *NSE4* was also mutated via error-prone PCR, and an n*se4-RAMP1* mutant containing three amino-acid changes (F311L, N324K, S329G) was found to produce a similar reduction in DSBs as *smc5* or *smc6* mutants (Figure 3C). These findings suggest that the effect on palindrome instability depends on the integrity of the Smc5/6 holoenzyme and on Smc5 interaction with Nse4.

**Figure 3.**
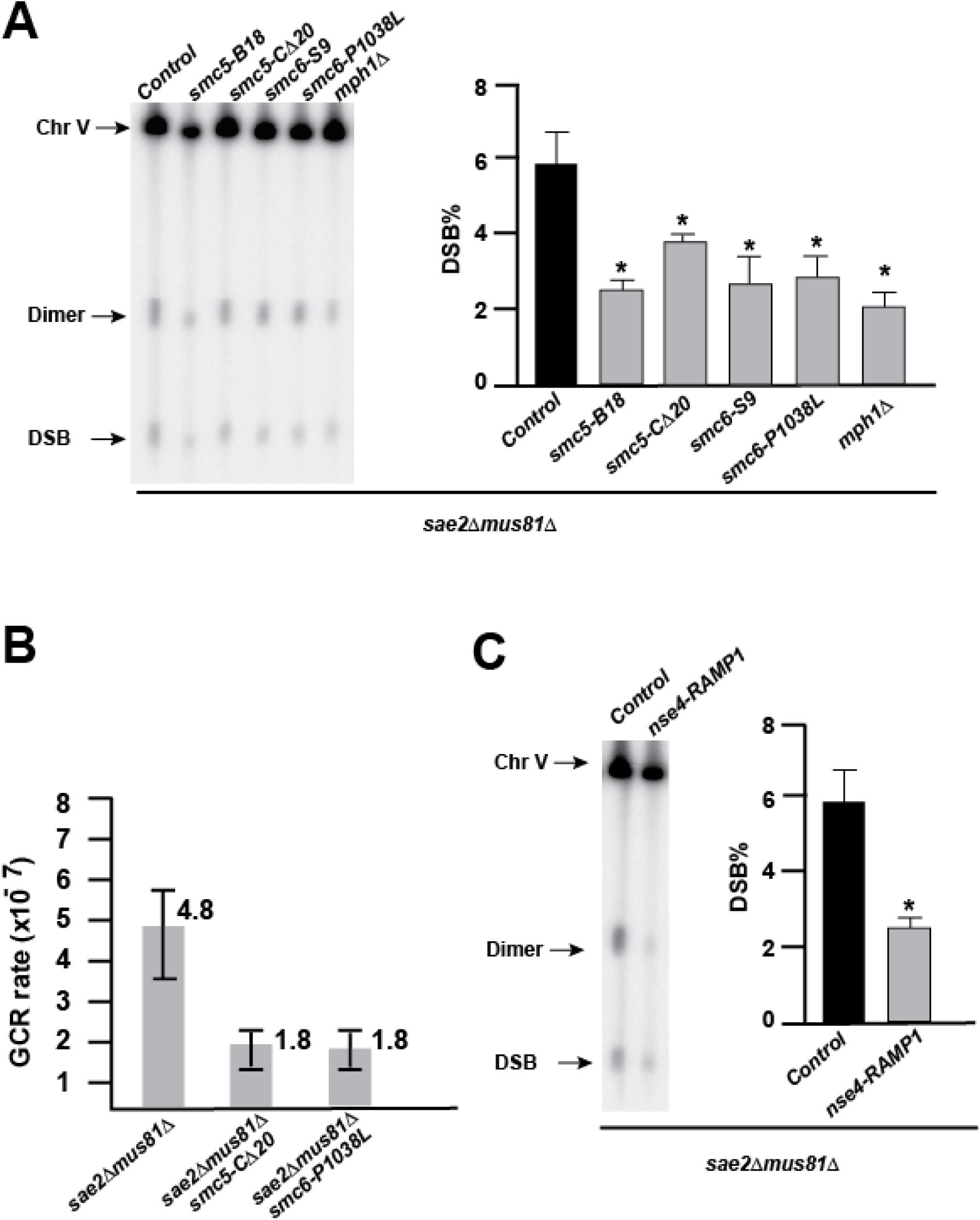
*smc5-C20*Δ*, smc6-P1038L* and *nse4-RAMP1* mutations reduce palindrome-mediated DSBs. A. Representative image of palindrome-mediated DSB detection by Southern blot of cells containing the indicated *smc5/6* mutations in a *sae2*Δ *mus81*Δ genetic background. Quantifications of DSBs are shown next to the gel image. Values are the mean of at least 3 independent biological replicates ± standard deviation; * *p* < 0.05. *P-*values were calculated by comparing mutants to their matched genetic control. B. *smc5-C20*Δ and *smc6-P1038L* affect *Alu*-PAL-induced GCR levels. Data are represented as the median value ± 95% confidence interval. C. Representative image of palindrome-mediated DSB detection by Southern blot of cells containing mutagenized *NSE4.* Quantifications of DSBs are shown next to the gel image. Values are the mean of at least three independent biological replicates ± standard deviation; * *p* < 0.05. *p* values were calculated by comparing mutants to their matched genetic control.

### Rad51, Rad54, Rtt107, and Smc5/6 are in the same pathway for palindrome regulation

To determine whether Mph1, Rad51, Rad54, Rtt107, and Smc5/6 are part of the same pathway involved in palindrome-mediated instability, multi-gene knockouts were created in a *sae2*Δ *mus81*Δ background, and DSBs were measured. Neither knockout nor mutation of Rad51, Rad54, Smc5/6, or Rtt107 caused a significant reduction in DSBs in a *mph1*Δ background (Figure 4A). Furthermore, the *smc5-C20*Δ and *smc6-P1038L* mutations did not decrease DSBs in *rad51*Δ (Figure 4 B). This indicates that Mph1, Rad51, Rad54, Rtt107, and Smc5/6 form an epistasis group. Overall, these findings suggest that Mph1, the Smc5/6 complex, Rad51, Rad54, and Rtt107 comprise a distinctive pathway that mediates palindrome instability.

**Figure 4.**
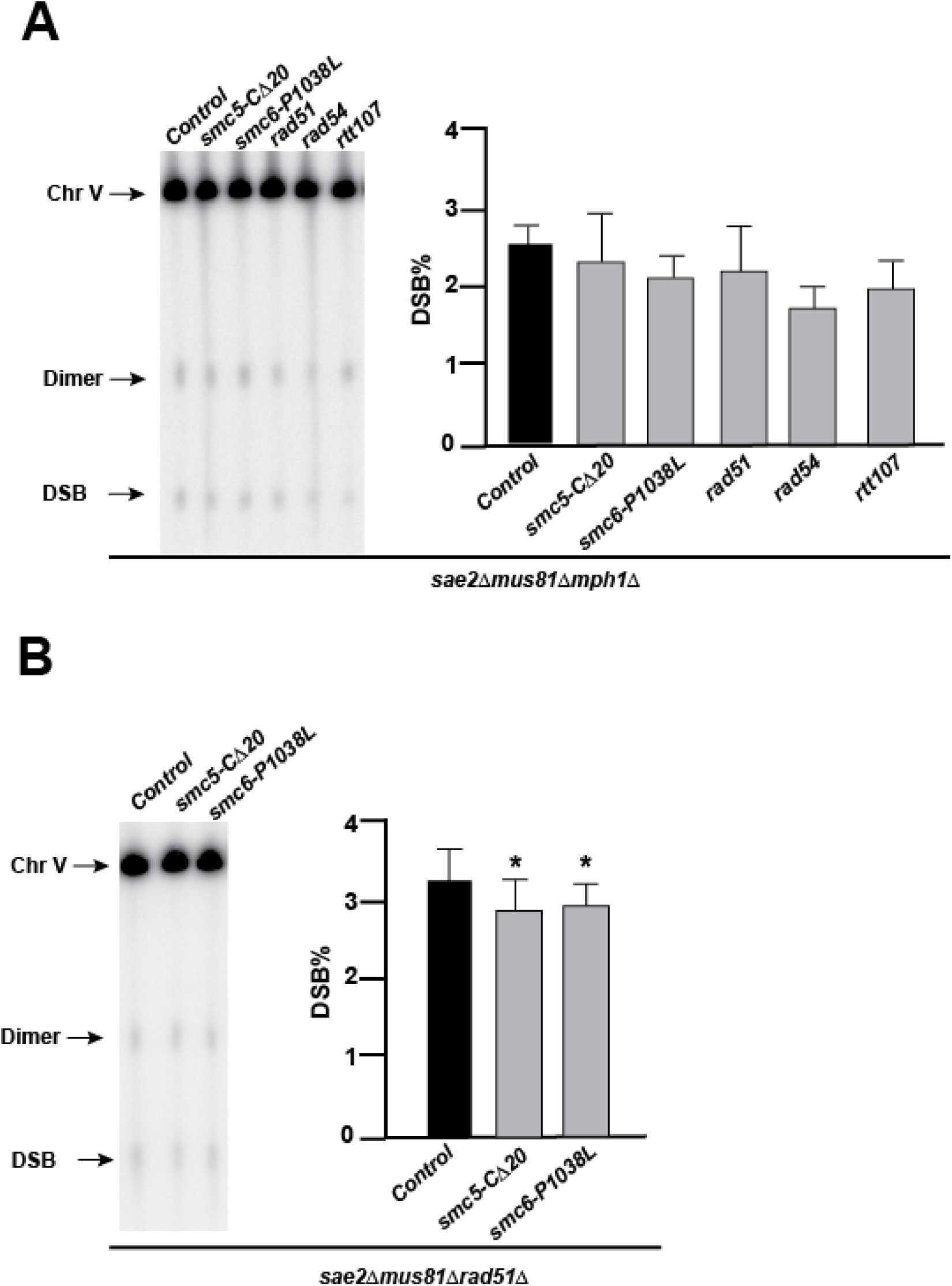
Mph1, Smc5/6, Rad51, Rad54, and Rtt107 form an epistasis group. A. Representative image of palindrome-mediated DSB detection by Southern blot of cells containing the indicated mutations or deletions in a *sae2*Δ *mus81*Δ *mph1*Δ genetic background. Quantifications of DSBs are shown next to the gel image. Values are the mean of at least 3 independent biological replicates ± standard deviation; * *p* < 0.05. *p* values were calculated by comparing mutants to their matched genetic control. B. Representative image of palindrome-mediated DSB detection by Southern blot of cells containing the indicated mutations or deletions in a *sae2*Δ *mus81*Δ *rad51*Δ background. Values are the mean of at least 3 independent biological replicates ± standard deviation; * *p* < 0.05. *p* values were calculated by comparing mutants to their matched genetic control.

### Mph1, but not Smc5/6, promotes the formation of cruciform structures at the replication fork

None of the proteins identified in the Mph1 pathway of palindrome-mediated instability possesses nuclease activity and therefore cannot be involved in resolving secondary structures. We thus hypothesized that the newly identified pathway affects palindrome fragility by promoting the formation of secondary structures, which, in turn, increases their susceptibility to nuclease cleavage. To determine whether Mph1 or Smc5/6 can promote secondary structure formation at replication forks, replication intermediate analysis by 2D DNA gel electrophoresis was performed using *sae2*Δ*mus81*Δ, *sae2*Δ*mus81*Δ*mph1*Δ and *sae2*Δ*mus81*Δ*smc6P1038L* mutant strains (Figure 5). Cells were arrested in G1 and then synchronously released into S phase. DNA was collected from the cells 50 minutes after release, when the replication fork is expected to traverse the region containing the palindrome, according to previous studies.

**Figure 5.**
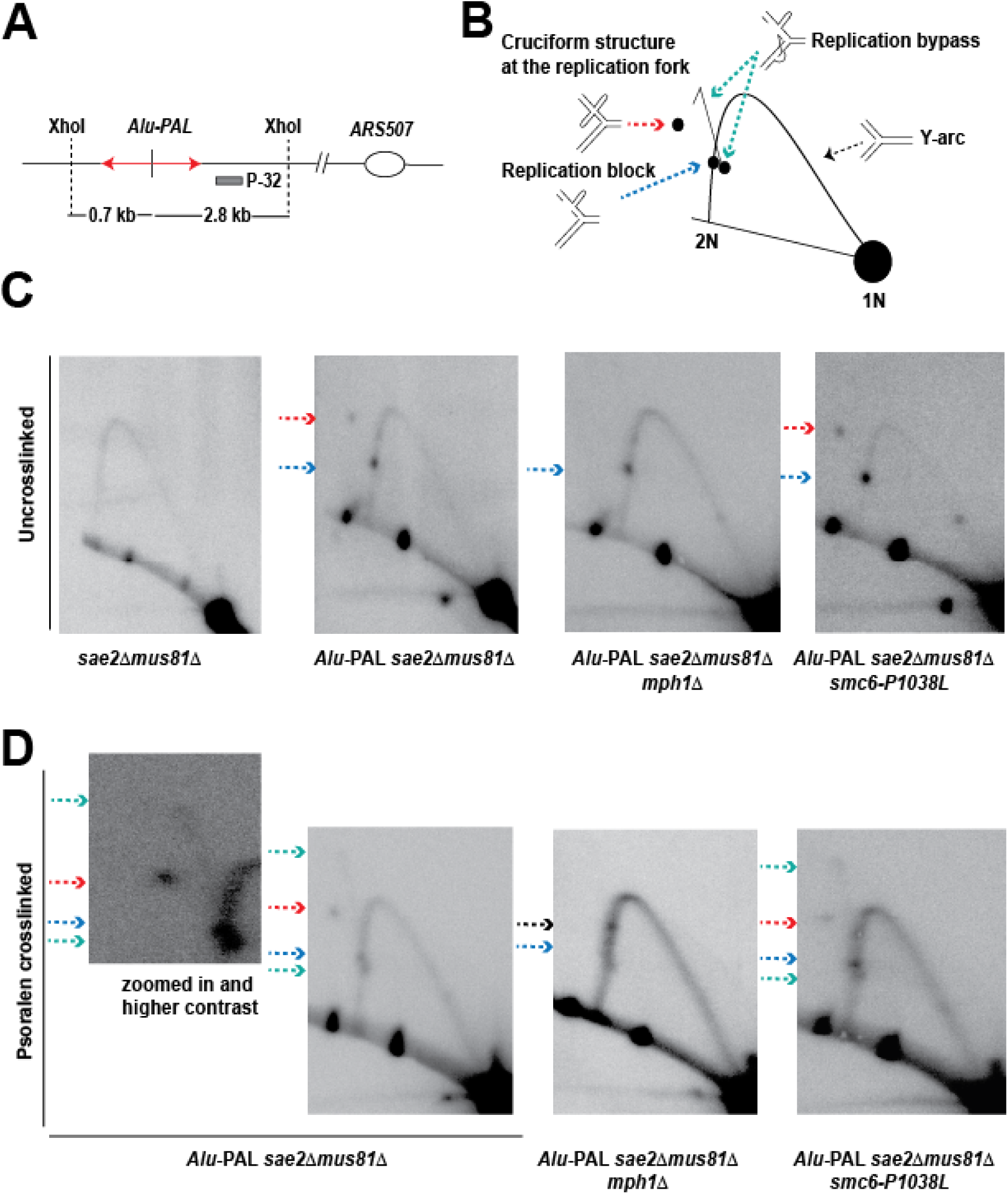
Mph1 promotes template switch-mediated cruciform formation. A. Illustration of restriction digestion with *Xho*I and the position of the probe used to reveal replication and template switch intermediates (grey rectangle). B. Scheme of replication and template switch intermediates. C. 2D gels analysis of not crosslinked replication and template switch intermediates. Left panel is a 2D analysis of the control strain that does not have the insertion of *Alu*-Pal in *LYS2*. Blue dotted arrow indicates replication block. Red dotted arrow shows cruciform structure formed at they replication fork. D. 2D gels analysis of psoralen- and UV- crosslinked replication and template switch intermediates. Green dotted arrows: deduced D-loop and D-loops with extended synthesis of the invaded strand. Black dotted arrow: fork reversal

Using 2D gel electrophoresis and placing the repeats on the long shoulder of the Y-arc, we previously showed that the hairpin structure formed by *Alu*-Pal causes replication fork pausing (18). In the current study, we revised the 2D analysis scheme and positioned the repeats on the short shoulder of the Y-arc (Figure 5A). In addition to the replication fork pause intermediates (indicated by blue arrows), which are specific to strains containing *Alu*-Pal, we also observed X-shaped intermediates (indicated by red arrows). These species migrated more slowly than the arrested forks along the Y-arc, indicating increased DNA mass and complexity, and likely represent cruciform structures at replication forks (Figure 5C). Notably, these structures were not visible in the *mph1*Δ mutant but were present in *smc6-P1038L*. The replication fork block remained unaffected by either Mph1 or Smc5/6 deficiency.

We used psoralen crosslinking to preserve secondary structures and prevent branch migration. In samples collected from cells containing Mph1, we detected additional intermediates (Figure 5D). An intermediate near the replication block that migrated faster suggests that these molecules lost base pairing at the sites where ethidium bromide cannot intercalate. Ethidium bromide is used in 2D gels to intercalate branched molecules and delay their movement in the gel. These structures most likely represent a template switch event involving the invasion of the 3’ sDNA strand from the lagging to the leading template and the formation of a D-loop (green arrow). The emerging spike (highlighted in the zoomed-in panel) consists of molecules with increased molecular mass, likely resulting from DNA synthesis following invasion (green arrows). These intermediates were also eliminated in *mph1*Δ. Interestingly, psoralen crosslinking in the *mph1*Δ mutant highlighted the accumulation of molecules present on the arc, which migrated with a delay relative to the replication block (black arrow). It is an expected pattern of migration for fork-reversal molecules, occurring when replication bypass in *mph1*Δ is unavailable. Overall, these data demonstrate that Mph1, but not Smc5/6, activity is required to promote the formation of a cruciform structure at replication forks during lesion bypass.

## Discussion

In this study, we uncovered the involvement of Mph1, Rad51, Rad54, Rtt107, and Smc5/6 in promoting DSB formation at the *Alu* palindrome in yeast. We discovered a pathway for cruciform formation involving replication bypass of the hairpin block mediated by Mph1, Rad51, and Rad54 proteins. We also propose a potential role for Rtt107 and Smc5/6 in facilitating the cleavage of the cruciform structure. A model for DSB formation at perfect palindromes and involving these proteins is presented in Figure 6.

**Figure 6.**
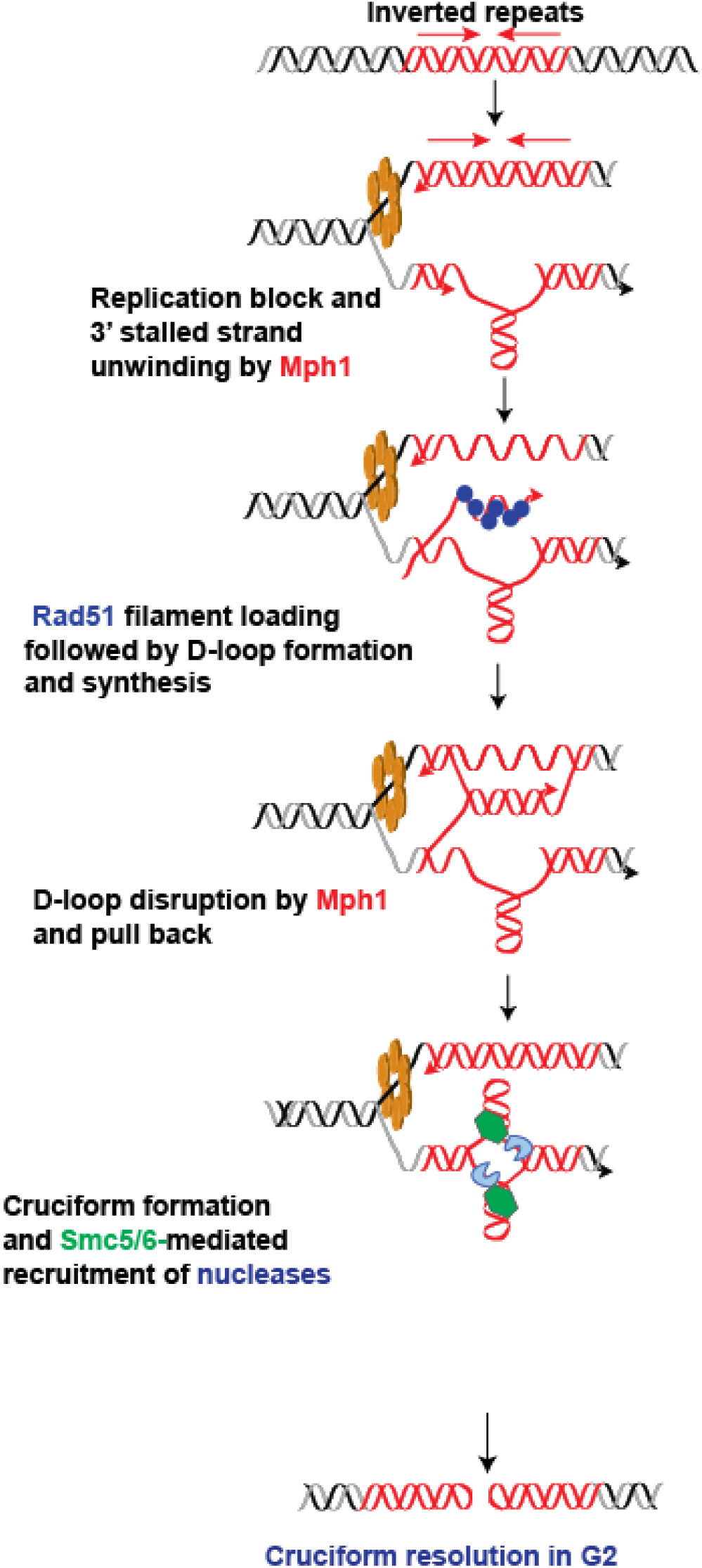
Model for cruciform formation resulting from replication template switch. See Discussion for details.

### The role of replication fork remodelling proteins Mph1, Rad51, and Rad54 in cruciform formation

Previously, we found that cruciforms can form through two pathways: one that relies on DNA synthesis and another that occurs independently of replication in G2 phase cells. Also, perfect palindromes and quasi-palindromes with spacers smaller than 9 bp effectively block replication by forming hairpins on the lagging strand (18). Based on our findings that deleting Rad51, Rad54, Mph1, or inactivating Mph1 helicase activity reduces palindrome-mediated DSBs and their known roles in strand remodeling during recombinational repair, we suggest that these components of the homologous recombination machinery are involved in a replication-dependent process of cruciform formation, which includes template switching when the replication fork encounters the hairpin structure (Figure 6). Mph1 is a 3’ to 5’ helicase (31) that can unwind the stalled newly synthesized strand at this replication barrier, creating a substrate for Rad51 filament loading. Rad51-mediated invasion of the stalled strand into the sister chromatid, aided by Rad54 (40), forms a D-loop, allowing DNA polymerase to extend the 3’ end of the stalled strand and synthesize the hairpin-forming sequence. Mph1 may also participate in the next step, given its known role in dismantling D-loops during homologous recombination (41). Dissolving the D-loop and reannealing the newly synthesized DNA might lead to cruciform structure formation.

Several lines of evidence support this cruciform formation mechanism. First, as shown in this study, intermediates corresponding to D-loop migration and cruciform-containing replication forks can be detected on 2D gels in strains containing Mph1, but are absent in *mph1* mutants (Figure 5). Second, Mph1’s dual role in D-loop formation and disruption has been linked to error-free bypass of DNA damage, involving sister-chromatid interactions. Mph1 works with Rad51 and Rad54 in this lesion bypass pathway (42,43). Third, in support of the proposed role for Mph1 and Rad51, the accumulation of X-shaped recombination intermediates in *esc2* and *smc5/6* mutants resulting from MMS-induced replication lesions is prevented in *mph1* and *rad51* mutants (34,44).

Overall, in this study, we deciphered one of the two pathways for cruciform formation, which involves the template switching mechanism between sister chromatids. Proteins that drive cruciform formation outside S-phase in G2 (18) still need to be identified.

### Rtt107 and Smc5/6 complex are mediators of palindrome fragility

We found that deletion of *RTT107* caused a reduction in palindrome-mediated DSBs (Figure 2A). We also isolated *smc5* and *smc6* mutants that exhibited reductions in DSBs in a *sae2*Δ *mus81*Δ, similar to the effect seen with *MPH1* deletion (Figure 2B). *smc5/6* mutants with only single mutations, *smc5-C20*Δ and *smc6-P1038L,* were sufficient to reduce palindrome-mediated DSB levels (Figure 3). Smc5/6, along with six non-SMC subunits (Nse1-6), form the Smc5/6 holo-complex (45). Structural analysis suggests that both Smc5-C20Δ and Smc6-P1038L lack regions that may interact with Nse4. Smc5-C20Δ lacks a domain within its head region that is inserted into the Nse4 wing-helix domain and likely promotes the interaction between the two subunits. The P1038 residue of Smc6 appears necessary to induce a turn of the peptide chain close to its key ATPase motifs. In addition, regions containing P1038 are also located close to Nse4 N-terminal helix-turn-helix region (46) and Supplemental Figure 2). We further provided evidence that Nse4 activity affects palindrome instability, as an *nse4-RAMP1* mutant exhibits DSB reduction similar to *smc5* and *smc6* mutants (Figure 3C). These data suggest that Nse4 and its interactions with Smc5 and Smc6 can be important for regulating palindrome breaks.

In *S. cerevisiae,* Smc5/6 associates with collapsed replication forks and facilitates their repair (47). Smc5/6 and Mph1 are known to interact and to be involved in creating and processing replication intermediates. The accumulation of toxic recombination intermediates in Smc5/6 mutants can be suppressed by the inactivation of Mph1 (34), suggesting that the Smc5/6 complex resolves or prevents toxic recombination intermediates promoted by Mph1. Smc5/6 can also restrain Mph1-mediated replication fork regression by binding to a region of Mph1 required for efficient regression. In contrast, Mph1-mediated D-loop dissociation is not regulated by Smc5/6 (37).

Based on these observations, we hypothesize that Smc5/6 promotes palindrome-mediated DSBs either by regulating replication fork remodelling and thus secondary structure formation or by mediating the resolution of cruciform structures generated by the homologous recombination machinery. We favor the second hypothesis because the pattern of intermediates corresponding to D-loop migration and cruciform-containing replication forks observed on 2D gels is identical in *SMC6* and *smc6-P1038L* strains (Figure 5C, D). Rtt107 has been identified as a hub that integrates multiple replication fork protection mechanisms, including the recruitment of Smc5/6 (35), the Slx1-Slx4 nuclease (48), and the activation of the Mus81-Mms4 nuclease (49). Smc5/6 also regulates Mus81-Mms4 activity in joint-molecule resolution and localization during meiosis (50). Therefore, we suggest that Rtt107 and Smc5/6 act together to recruit a nuclease that cleaves the cruciform. Although we found that neither Slx1-Slx4 nor Mus81-Mms4 structure-specific nucleases are involved in cruciform resolution at *Alu*-Pal (18), it is possible that another cruciform-cutting enzyme can be recruited and/or activated by Rtt107 and Smc5/6. It has been found that Smc5/6 stably binds to synthetic replication fork and joint molecules consisting of stretches of dsDNA and ssDNA (51), suggesting that the cruciform structure can be a potential substrate for Smc5/6 binding. Future studies testing this model can provide insights into the roles of the uncovered proteins in palindrome instability.

## Materials and Methods

### Yeast strains and plasmids

The strain used for the genome-wide screen to identify mutants with low *Alu* palindrome fragility, which contains GCR and gene amplification cassettes on chromosome VI, is YKL4143 and has the following genotype: MATa, *bar1*Δ*, trp1*Δ*, his3*Δ*, ura3*Δ*, leu2*Δ*, ade2*Δ*, lys2*Δ*, sfa1*Δ, *yhr054c*Δ *cup2*Δ, *VI* 253403:: *lys2*::*Alu*-PAL *SFA1CUP1phMX ADE2,CAN1*, *mus81::natMX,, sae2::TRP1*.

The strains used in this study to determine the effect of mutations in Mph1, Rad51, Rad54, Rtt1107, Smc5 Smc6, and Nse4 genes on GCRs, DSB levels and 2D patterns are isogenic and are built on YKL719 with the following genotype: a *MAT*α*, bar1*Δ, *his7–2, trp1*Δ, *ura3*Δ, *leu2–3,112, ade2*Δ, *lys2*Δ, *cup1*Δ *yhr054c*Δ *cup2*Δ, *V34205::ADE2 lys2::Alu-Pal V29616::CUP1* background. The gross chromosomal rearrangement (GCR) cassette and palindromic sequence insertion are described in Ait Saada et al., 2021. For all experiments, yeast strains were freshly thawed from glycerol stocks. All strains were grown at 30°C and manipulated using standard techniques. Unless otherwise specified, the medium used was YPD (1% yeast extract, 2% peptone, and 2% dextrose). Gene knockouts and introduction of mutant alleles were conducted with one of the following antibiotic resistance or prototrophic markers: *kanMX*, *hphMX*, *natMX*, *TRP1* or *URA3*.

Genomic fragments containing *SMC5*, *SMC6*, and *NSE4* genes were cloned into the polylinker of pRS414 between PstI and KpnI sites, and *URA3* was inserted approximately 200 bp downstream of each ORF via overlap extension PCR (52). This process resulted in the creation of pKL685, pKL686, and pKL703 vectors, respectively. These vectors were used to generate libraries of randomly mutagenized *SMC5-URA3*, *SMC6-URA3*, and *NSE4-URA3* cassettes. To introduce random mutations, standard PCR reactions were performed with ExTaq polymerase (Takara), using MnSO_4_ at a final concentration of 640 μM. The YKL4143 strain was transformed with the mutagenized libraries, and Ura+ colonies were screened for hypo-fragility mutants. pKL685 and pKL686 were used to generate individual mutations from *smc5-B18* and *smc6-S9* alleles via site-directed mutagenesis. These plasmids, including those carrying *smc5-C20*Δ (pKL712) and *smc6-P1038L* (pKL705) alleles responsible for the hypo-fragility effect, were used to replace *SMC5* and *SMC6* chromosomal alleles. The *Mph1* genomic fragment was cloned into the polylinker of pUC19 between KpnI and BamHI sites, with *kanMX* inserted approximately 250 bp downstream of the *MPH1* ORF, generating the pKL666 vector. This vector was used to create *mph1-DEAH*Δ, *mph1-Q603D*, and *mph1-S98P* alleles via site-directed mutagenesis, resulting in pKL667, pKL668, and pKL669 vectors, respectively. These plasmids were used to replace the wild-type *MPH1* copy with the *mph1-S98P-kanMX*, *mph1-DEAH*Δ*-kanMX*, and *mph1-Q603D-kanMX* cassettes. Oligonucleotide sequences used for random and site-directed mutagenesis and for overlap extension PCR are available upon request.

### Genome-wide screen for mutants displaying diminished palindrome-induced instability

YKL4143 cells were treated with ethyl methanesulfonate (EMS, 1 μl/1 ml of saturated yeast culture) to achieve approximately 5 mutations per cell, simplifying downstream analysis. Mutants were plated on YPD to produce single colonies, which were then replica-plated onto indicative media to screen for reduced palindrome-mediated GCRs and gene amplification compared to the nonmutated parental strain. GCRs were identified by the appearance of red, canavanine-resistant colonies on canavanine-containing plates [arginine-dropout medium containing 4 mg/l adenine and 60 mg/l L-canavanine]. Gene amplification of *CUP1* and *SFA1*, conferring drug resistance, was assessed on synthetic media with copper (200 µM) or formaldehyde (3 mM), respectively (Supplemental Figure 1). Mutants were screened on YPG (1% yeast extract, 2% peptone, 3% glycerol) and arginine-dropout media to exclude petite mutants and arginine auxotrophs, respectively. For mutants showing both decreased GCRs and gene amplification, double-strand break (DSB) detection was performed using clamped homogeneous electric field (CHEF) electrophoresis and Southern blotting, as described in the “DSB detection” section. Approximately 113,000 colonies were screened, leading to the identification of two mutants with diminished palindrome-mediated DSBs.

### Whole genome sequencing

Mutants that displayed significantly reduced DSB levels compared to the parental strain were subjected to whole genome sequencing performed by Psomagen (Rockville, MD) using an Illumina TruSeq DNA Nano 350 bp library prep and an Illumina NovaSeq 6000. Reads were aligned to the *S. cerevisiae* reference genome (SacCer3) using the Burrows-Wheeler Aligner (53), Picard (http://broadinstitute.github.io/picard), SAMtools (54), and Genome Analysis Toolkit v3.8 (55). Variants were called against the non-mutated parental strain using Mutect2 (55).

### GCR rate calculation

GCR rates were calculated by the fluctuation test. Strains were grown on solid YPD medium for 3 days. For each strain, 12-16 independent colonies were selected for a fluctuation test (56). Appropriate dilutions of cells were plated on YPD and canavanine plates [arginine-drop out medium containing adenine (4 mg/l) and L-canavanine (60 mg/l)]. Red colonies on canavanine media reflect GCRs and white colonies reflect loss of *CAN1* activity through sporadic mutations. GCR and mutation rates were calculated as previously described (57).

### DSB detection

For DSB detection, cells were grown overnight but not to saturation. Approximately 4 × 10^8^ cells per sample were embedded in a 0.75% low-melting-point agarose plug. Each plug was treated with 0.5 mg of zymolyase at 30°C for about 6 hours and 1 mg of proteinase K at 30°C overnight. For each sample, roughly a quarter of a plug (around 40 μl) was cut out and equilibrated in electrophoresis running buffer (0.5x TBE) for 20 minutes. The sample was then cast into a 1% pulsed-field gel electrophoresis (PFGE) certified agarose gel and separated by contour-clamped homogeneous electric field (CHEF) electrophoresis. To distinguish the DSB fragment (∼29 kb for chromosome VI and ∼43 kb for chromosome V) from intact chromosome (∼ 270 kb chromosome VI and ∼580 kb chromosome V), samples were run for 26 hours at 6 V/cm and 14°C with an included angle of 120°, an initial switching time of 3.14 seconds, and a final switching time of 7.68 seconds. Gels containing separated DNA fragments were treated sequentially with 0.25 N HCl (25 minutes), an alkaline buffer (1.5 M NaCl, 0.5 M NaOH, 30 minutes), and a neutralization buffer (1.5 M NaCl, 1 M Tris, pH 7.5, 30 minutes). DNA was then transferred to a charged nylon membrane in 10x SSC for 2 hours at 70 mmHg using Posiblotter (Stratagene). Southern hybridization was performed overnight with a P^32^-radiolabeled *LYS2-* (for chromosome VI) or *HPA3*-specific probe (for chromosome V) in PerfectHyb Plus Hybridization Buffer (MilliporeSigma) at 68°C. Membranes were washed twice in 0.1× SSC with 0.1% SDS at 68°C and then exposed to a phosphor storage screen. The bands were visualized by phosphorimaging with a Typhoon imager (GE Healthcare), and densitometry analysis of the broken fragments was conducted using ImageJ software (National Institutes of Health).

### Replication intermediate analysis by 2D gel electrophoresis

Strains were cultured overnight in 400 ml of YPD to an OD600 of 0.8. Alpha-factor (0.1 μg/ml) was added to the media to arrest cells in G1 and cells were cultured for an additional 3 h. Cells were washed twice with an equivalent volume of water and resuspended in 800 ml of YPD with pronase E (12.5 μg/ml) to release cells into S phase. Fifty minutes after release, each culture was collected in the presence of frozen EDTA (pH 8, 20 mM final) and sodium azide (0.1% final). Cells were disrupted by grinding in liquid nitrogen using the Spex Freezer Mill 6870 and genomic DNA was extracted by cesium chloride gradient as described in https://fangman-brewer.genetics.washington.edu/DNA_prep.html. DNA was digested with XhoI (50 units) for several hours and precipitated in ethanol. To cut cruciform-containing molecules, genomic DNA after XhoI digestion was treated with 5 μg of RuvC (Diagno Cine) for 30 minutes. Restriction digestion fragments were separated in a 0.4% agarose gel (1x TBE at 1.7 V/cm for 22 h). Gel slices containing the fragment of interest were excised and cast into a 1.2% agarose gel made with running buffer and run in 1x TBE with ethidium bromide (3 mg/ml) at 6 V/cm for 10 h at 4°C. Gel processing and Southern hybridization were performed as described above. Replication intermediates were highlighted with a *LYS2*-specific probe.

*In vivo* psoralen crosslinking was used to stabilize DNA intermediates that are prone to destruction during the DNA isolation protocol. ∼ 5x 10^9^ cells were collected after release from alpha factor, as described above, and resuspended in 3 ml of cold water. The cell suspensions were transferred to petri dishes kept on ice, and 300 μL of 4,5’,8-trimethylpsoralen (0.2 mg/mL in 10% ethanol) was added and mixed carefully. After incubation in the dark for 5 minutes, the cell suspensions were irradiated for 10 minutes at 365 nm using a Stratagene UV Stratalinker 2400. This treatment was repeated twice more. Genomic DNA was extracted as described above. Gels after resolving psoralen-crosslinked replication intermediates in the second dimension were irradiated at 254 nm with a dose of 5.55 kJ/m² (5550 energy setting on the Stratagene UV Stratalinker 2400) to decrosslink the DNA samples and facilitate transfer and hybridization with a probe.

### Statistical analysis

For DSB detection, all data presented are averages of at least 3 biological replicates. DSB percentage corresponds to the proportion of the DSB fragment relative to the total signal of chromosome V (unbroken chromosome V + DSB). Average values with error bars indicating standard deviation are shown. Data were analyzed and visualized with Prism9 (GraphPad). Statistical comparisons were performed using *t*-tests (to compare two samples) or ANOVA with a Dunnett’s Multiple Comparison (to compare more than two samples). Equal variances were tested with a Brown-Forsyth Test.

### Sequence alignment and structure analysis

Sequences were obtained from UniProt (58), and multiple sequence alignment was performed using Clustal Omega(59) with default parameters. Alignment results were visualized using the ESPript3 online tool (60). Analyses of the surface were performed using the default settings in ChimeraX (61).

---

**Supplemental Figure 1.** Genome-wide screen identifies Mph1 as a contributor to palindrome instability. A. GCR and gene amplification assays on the right arm of chromosome VI to monitor *Alu-*PAL-induced instability. The *LYS2* probe (grey rectangle) is used to highlight DSBs, dimers, and intact chromosome VI. Lower panels: examples of analysis of ethyl methanesulfonate-mutated cells replica-plated on media indicative of gene amplification (synthetic media with 200 μM copper and 3 mM formaldehyde) and gross chromosomal rearrangements (canavanine). A patch corresponding to the non-mutated control strain is indicated with squares, while a patch for the *mph1* mutant is marked with circles. B. Detection of palindrome-mediated DSBs by Southern blot of cells with the indicated deletions. An isolate exhibiting low GCR and gene amplification frequency was sequenced and found to have point mutations in the *MPH1*, *ATG1*, *PCA1*, and *YKL225W* genes. Individual disruptions were created in the YKL4143 strain, which contains *sae2*Δ*mus81*Δ disruptions, to identify mutants responsible for the hypo-fragility phenotype. The diagram indicates a 29 kb broken fragment, a 58 kb inverted dimer resulting from replication of hairpin-capped breaks, and intact chromosome VI. C. Effect of *mph1*Δ on Alu-induced GCR and gene amplification using chromosome VI assays. Gene amplification rates were measured with media containing 200 μM copper and 3 mM formaldehyde.

**Supplemental Figure 2.** Structure and sequence analysis of amino acid changes in *smc5-A6*, *smc5-B18* and *smc6-S9* alleles. A. Summary of the features of the mutated residues. The locations of the mutated residues within the head, arm, and hinge regions of the proteins and secondary structures containing the mutated residues are indicated. Sequence similarities were assessed by comparing Smc5 and Smc6 homologs across seven species, including *Saccharomyces cerevisiae*, *Schizosaccharomyces pombe*, *Caenorhabditis elegans*, *Drosophila melanogaster*, *Xenopus laevis*, *Mus musculus*, and *Homo sapiens*. High similarity indicates identical or similar residues among homologs, and low similarity indicates that residues with different biochemical properties are present in the sequence alignment. The last column includes residues found in six related yeast species, including *S. arboricola*, *S. boulardii*, *S. eubayanus*, *S. mikatae*, *S. paradoxus*, and *S. pastorianus*. B. The C-terminal 20 amino acids of Smc5 directly contact the Nse4 protein based on the cryo-EM structure of the yeast Smc5/6 complex (PDB: 7TVE). C. Multiple sequence alignment of the Smc5 C-terminal region containing the 20 residues deleted in the *smc5-B18* allele (ΔC20). S. cer (*Saccharomyces cerevisiae*), S. pom (*Schizosaccharomyces pombe*), C. ele (*Caenorhabditis elegans*), D. mel (*Drosophila melanogaster*), X. lae (*Xenopus laevis*), M. mus (*Mus musculus*), and H. sap (*Homo sapiens*). Similar residues are shown in red and low similarity residues in black. D. Smc6-P1038 is essential for the correct positioning of the ATPase Walker B motif and the C-loop of the ATPase domain based on the cryo-EM structure of the yeast Smc5/6 complex (PDB: 7TVE). The rigidity of the P1038 side chain ensures proper separation and function of these two motifs. E. Multiple sequence alignments show that while the yeast Smc6 protein contains P1038 to allow the turning of the peptide chain, Smc6 homologs from other species contain a nearby proline that may achieve a similar function. The species shown here and the coloring scheme are the same as in A-C.

## Acknowledgement

We are very thankful to Dr. You Yu for the initial structural analyses. Funding of this work was provided by NIH grant R01GM158860 (to K.S.L.) and R35GM14526 (to X.Z.). J. F. is supported by the Postdoctoral Fellowship grant PF-24-1318483-01-DMC from the American Cancer Society

## Author contributions

Conceptualization: A.B.C., A.A.S. and K.S.L.; Methodology: A.B.C., A.A.S., X.Z. and K.S.L.; Investigation: A.A.S., A.B.C., R.H., J.F. and T.P.; Writing – Original Draft: A.B.C. and K.S.L.; Writing – Review & Editing: A.B.C, A.A.S., J.F. X.Z. and K.S.L.; Funding Acquisition: K.S.L. and X.Z.; Resources: K.S.L.; Supervision: K.S.L.; Project Administration: K.S.L.

## Declaration on Interests

The authors declare no competing interests.

## Notes

### Competing Interest Statement

The authors have declared no competing interest.

## References

1. Ganapathiraju, M.K., Subramanian, S., Chaparala, S. and Karunakaran, K.B. (2020) A reference catalog of DNA palindromes in the human genome and their variations in 1000 Genomes. Human genome variation, 7, 1–12.

2. Guenthoer, J., Diede, S.J., Tanaka, H., Chai, X., Hsu, L., Tapscott, S.J. and Porter, P.L. (2012) Assessment of palindromes as platforms for DNA amplification in breast cancer. Genome Res, 22, 232–245.

3. Kato, T., Franconi, C.P., Sheridan, M.B., Hacker, A.M., Inagakai, H., Glover, T.W., Arlt, M.F., Drabkin, H.A., Gemmill, R.M., Kurahashi, H. et al. (2014) Analysis of the t(3;8) of hereditary renal cell carcinoma: a palindrome-mediated translocation. Cancer Genet, 207, 133–140.

4. Kato, T., Kurahashi, H. and Emanuel, B.S. (2012) Chromosomal translocations and palindromic AT-rich repeats. Curr Opin Genet Dev, 22, 221–228.

5. Mangano, R., Piddini, E., Carramusa, L., Duhig, T., Feo, S. and Fried, M. (1998) Chimeric amplicons containing the c-myc gene in HL60 cells. Oncogene, 17, 2771–2777.

6. Neiman, P.E., Elsaesser, K., Loring, G. and Kimmel, R. (2008) Myc oncogene-induced genomic instability: DNA palindromes in bursal lymphomagenesis. PLoS Genet, 4, e1000132.

7. Neiman, P.E., Kimmel, R., Icreverzi, A., Elsaesser, K., Bowers, S.J., Burnside, J. and Delrow, J. (2006) Genomic instability during Myc-induced lymphomagenesis in the bursa of Fabricius. Oncogene, 25, 6325–6335.

8. Rooks, H., Clark, B., Best, S., Rushton, P., Oakley, M., Thein, O.S., Cuthbert, A.C., Britland, A., Ruf, A. and Thein, S.L. (2012) A novel 506kb deletion causing εγδβ thalassemia. Blood Cells Mol Dis, 49, 121–127.

9. Tanaka, H., Bergstrom, D.A., Yao, M.C. and Tapscott, S.J. (2005) Widespread and nonrandom distribution of DNA palindromes in cancer cells provides a structural platform for subsequent gene amplification. Nat Genet, 37, 320–327.

10. Tanaka, H., Bergstrom, D.A., Yao, M.C. and Tapscott, S.J. (2006) Large DNA palindromes as a common form of structural chromosome aberrations in human cancers. Hum Cell, 19, 17–23.

11. Tanaka, H., Cao, Y., Bergstrom, D.A., Kooperberg, C., Tapscott, S.J. and Yao, M.C. (2007) Intrastrand annealing leads to the formation of a large DNA palindrome and determines the boundaries of genomic amplification in human cancer. Mol Cell Biol, 27, 1993–2002.

12. Zhu, H., Shang, D., Sun, M., Choi, S., Liu, Q., Hao, J., Figuera, L.E., Zhang, F., Choy, K.W., Ao, Y. et al. (2011) X-linked congenital hypertrichosis syndrome is associated with interchromosomal insertions mediated by a human-specific palindrome near SOX3. Am J Hum Genet, 88, 819–826.

13. Vlaar, J.M., Borgman, A., Kalkhoven, E., Westland, D., Besselink, N., Shale, C., Faltas, B.M., Priestley, P., Kuijk, E. and Cuppen, E. (2022) Recurrent exon-deleting activating mutations in AHR act as drivers of urinary tract cancer. Sci Rep, 12, 10081.

14. Ait Saada, A., Guo, W., Costa, A.B., Yang, J., Wang, J. and Lobachev, K.S. (2023) Widely spaced and divergent inverted repeats become a potent source of chromosomal rearrangements in long single-stranded DNA regions. Nucleic Acids Res, 51, 3722–3734.

15. Leach, D.R. (1994) Long DNA palindromes, cruciform structures, genetic instability and secondary structure repair. Bioessays, 16, 893–900.

16. Osia, B., Twarowski, J., Jackson, T., Lobachev, K., Liu, L. and Malkova, A. (2022) Migrating bubble synthesis promotes mutagenesis through lesions in its template. Nucleic Acids Res, 50, 6870–6889.

17. Eykelenboom, J.K., Blackwood, J.K., Okely, E. and Leach, D.R. (2008) SbcCD causes a double-strand break at a DNA palindrome in the Escherichia coli chromosome. Mol Cell, 29, 644–651.

18. Ait Saada, A., Costa, A.B., Sheng, Z., Guo, W., Haber, J.E. and Lobachev, K.S. (2021) Structural parameters of palindromic repeats determine the specificity of nuclease attack of secondary structures. Nucleic Acids Res, 49, 3932–3947.

19. Neelsen, K.J. and Lopes, M. (2015) Replication fork reversal in eukaryotes: from dead end to dynamic response. Nature reviews Molecular cell biology, 16, 207–220.

20. Quinet, A., Lemaçon, D. and Vindigni, A. (2017) Replication fork reversal: players and guardians. Molecular cell, 68, 830–833.

21. Giannattasio, M., Zwicky, K., Follonier, C., Foiani, M., Lopes, M. and Branzei, D. (2014) Visualization of recombination-mediated damage bypass by template switching. Nature structural & molecular biology, 21, 884–892.

22. Sinden, R.R., Zheng, G.X., Brankamp, R.G. and Allen, K.N. (1991) On the deletion of inverted repeated DNA in Escherichia coli: effects of length, thermal stability, and cruciform formation in vivo. Genetics, 129, 991–1005.

23. Havas, K., Flaus, A., Phelan, M., Kingston, R., Wade, P.A., Lilley, D.M. and Owen-Hughes, T. (2000) Generation of superhelical torsion by ATP-dependent chromatin remodeling activities. Cell, 103, 1133–1142.

24. Jaskelioff, M., Van Komen, S., Krebs, J.E., Sung, P. and Peterson, C.L. (2003) Rad54p is a chromatin remodeling enzyme required for heteroduplex DNA joint formation with chromatin. J Biol Chem, 278, 9212–9218.

25. Yu, S., Owen-Hughes, T., Friedberg, E.C., Waters, R. and Reed, S.H. (2004) The yeast Rad7/Rad16/Abf1 complex generates superhelical torsion in DNA that is required for nucleotide excision repair. DNA Repair (Amst), 3, 277–287.

26. Feng, X., Xie, F.Y., Ou, X.H. and Ma, J.Y. (2020) Cruciform DNA in mouse growing oocytes: Its dynamics and its relationship with DNA transcription. PLoS One, 15, e0240844.

27. Kouzine, F., Sanford, S., Elisha-Feil, Z. and Levens, D. (2008) The functional response of upstream DNA to dynamic supercoiling in vivo. Nat Struct Mol Biol, 15, 146–154.

28. Narayanan, V., Mieczkowski, P.A., Kim, H.M., Petes, T.D. and Lobachev, K.S. (2006) The pattern of gene amplification is determined by the chromosomal location of hairpin-capped breaks. Cell, 125, 1283–1296.

29. Liu, H., Krizek, J. and Bretscher, A. (1992) Construction of a GAL1-regulated yeast cDNA expression library and its application to the identification of genes whose overexpression causes lethality in yeast. Genetics, 132, 665–673.

30. Whitby, M.C. (2010) The FANCM family of DNA helicases/translocases. DNA Repair (Amst), 9, 224–236.

31. Prakash, R., Krejci, L., Van Komen, S., Anke Schurer, K., Kramer, W. and Sung, P. (2005) Saccharomyces cerevisiae MPH1 gene, required for homologous recombination-mediated mutation avoidance, encodes a 3’ to 5’ DNA helicase. J Biol Chem, 280, 7854–7860.

32. Scheller, J., Schürer, A., Rudolph, C., Hettwer, S. and Kramer, W. (2000) MPH1, a yeast gene encoding a DEAH protein, plays a role in protection of the genome from spontaneous and chemically induced damage. Genetics, 155, 1069–1081.

33. Xue, X., Sung, P. and Zhao, X. (2015) Functions and regulation of the multitasking FANCM family of DNA motor proteins. Genes Dev, 29, 1777–1788.

34. Chen, Y.-H., Choi, K., Szakal, B., Arenz, J., Duan, X., Ye, H., Branzei, D. and Zhao, X. (2009) Interplay between the Smc5/6 complex and the Mph1 helicase in recombinational repair. Proceedings of the National Academy of Sciences, 106, 21252–21257.

35. Hang, L.E., Peng, J., Tan, W., Szakal, B., Menolfi, D., Sheng, Z., Lobachev, K., Branzei, D., Feng, W. and Zhao, X. (2015) Rtt107 Is a Multi-functional Scaffold Supporting Replication Progression with Partner SUMO and Ubiquitin Ligases. Mol Cell, 60, 268–279.

36. Leung, G.P., Lee, L., Schmidt, T.I., Shirahige, K. and Kobor, M.S. (2011) Rtt107 is required for recruitment of the SMC5/6 complex to DNA double strand breaks. J Biol Chem, 286, 26250–26257.

37. Xue, X., Choi, K., Bonner, J.N., Chiba, T., Kwon, Y., Xu, Y., Sanchez, H., Wyman, C., Niu, H. and Zhao, X. (2014) Restriction of replication fork regression activities by a conserved SMC complex. Molecular cell, 56, 436–445.

38. Zapatka, M., Pocino-Merino, I., Heluani-Gahete, H., Bermudez-Lopez, M., Tarres, M., Ibars, E., Sole-Soler, R., Gutierrez-Escribano, P., Apostolova, S. and Casas, C. (2019) Sumoylation of Smc5 promotes error-free bypass at damaged replication forks. Cell Reports, 29, 3160–3172. e3164.

39. Palecek, J., Vidot, S., Feng, M., Doherty, A.J. and Lehmann, A.R. (2006) The Smc5-Smc6 DNA repair complex. bridging of the Smc5-Smc6 heads by the KLEISIN, Nse4, and non-Kleisin subunits. J Biol Chem, 281, 36952–36959.

40. Petukhova, G., Van Komen, S., Vergano, S., Klein, H. and Sung, P. (1999) Yeast Rad54 promotes Rad51-dependent homologous DNA pairing via ATP hydrolysis-driven change in DNA double helix conformation. J Biol Chem, 274, 29453–29462.

41. Prakash, R., Satory, D., Dray, E., Papusha, A., Scheller, J., Kramer, W., Krejci, L., Klein, H., Haber, J.E. and Sung, P. (2009) Yeast Mph1 helicase dissociates Rad51-made D-loops: implications for crossover control in mitotic recombination. Genes & development, 23, 67–79.

42. Ede, C., Rudolph, C.J., Lehmann, S., Schurer, K.A. and Kramer, W. (2011) Budding yeast Mph1 promotes sister chromatid interactions by a mechanism involving strand invasion. DNA Repair (Amst), 10, 45–55.

43. Panico, E.R., Ede, C., Schildmann, M., Schurer, K.A. and Kramer, W. (2010) Genetic evidence for a role of Saccharomyces cerevisiae Mph1 in recombinational DNA repair under replicative stress. Yeast, 27, 11–27.

44. Mankouri, H.W., Ngo, H.P. and Hickson, I.D. (2009) Esc2 and Sgs1 act in functionally distinct branches of the homologous recombination repair pathway in Saccharomyces cerevisiae. Mol Biol Cell, 20, 1683–1694.

45. Yu, Y., Li, S., Ser, Z., Sanyal, T., Choi, K., Wan, B., Kuang, H., Sali, A., Kentsis, A., Patel, D.J. et al. (2021) Integrative analysis reveals unique structural and functional features of the Smc5/6 complex. Proc Natl Acad Sci U S A, 118.

46. Jo, A., Li, S., Shin, J.W., Zhao, X. and Cho, Y. (2021) Structure basis for shaping the Nse4 protein by the Nse1 and Nse3 dimer within the Smc5/6 complex. Journal of molecular biology, 433, 166910.

47. Ampatzidou, E., Irmisch, A., O’Connell, M.J. and Murray, J.M. (2006) Smc5/6 is required for repair at collapsed replication forks. Mol Cell Biol, 26, 9387–9401.

48. Ohouo, P.Y., Bastos de Oliveira, F.M., Almeida, B.S. and Smolka, M.B. (2010) DNA damage signaling recruits the Rtt107-Slx4 scaffolds via Dpb11 to mediate replication stress response. Mol Cell, 39, 300–306.

49. Princz, L.N., Wild, P., Bittmann, J., Aguado, F.J., Blanco, M.G., Matos, J. and Pfander, B. (2017) Dbf4-dependent kinase and the Rtt107 scaffold promote Mus81-Mms4 resolvase activation during mitosis. EMBO J, 36, 664–678.

50. Copsey, A., Tang, S., Jordan, P.W., Blitzblau, H.G., Newcombe, S., Chan, A.C., Newnham, L., Li, Z., Gray, S., Herbert, A.D. et al. (2013) Smc5/6 coordinates formation and resolution of joint molecules with chromosome morphology to ensure meiotic divisions. PLoS Genet, 9, e1004071.

51. Chang, J.T., Li, S., Beckwitt, E.C., Than, T., Haluska, C., Chandanani, J., O’Donnell, M.E., Zhao, X. and Liu, S. (2022) Smc5/6’s multifaceted DNA binding capacities stabilize branched DNA structures. Nat Commun, 13, 7179.

52. Bryksin, A.V. and Matsumura, I. (2010) Overlap extension PCR cloning: a simple and reliable way to create recombinant plasmids. Biotechniques, 48, 463–465.

53. Li, H. and Durbin, R. (2009) Fast and accurate short read alignment with Burrows-Wheeler transform. Bioinformatics, 25, 1754–1760.

54. Li, H., Handsaker, B., Wysoker, A., Fennell, T., Ruan, J., Homer, N., Marth, G., Abecasis, G., Durbin, R. and Genome Project Data Processing, S. (2009) The Sequence Alignment/Map format and SAMtools. Bioinformatics, 25, 2078–2079.

55. Van der Auwera, G.A. and O’Connor, B.D. (2020) Genomics in the cloud: using Docker, GATK, and WDL in Terra. O’Reilly Media.

56. Drake, J.W. (1991) A constant rate of spontaneous mutation in DNA-based microbes. Proc Natl Acad Sci U S A, 88, 7160–7164.

57. Lobachev, K.S., Gordenin, D.A. and Resnick, M.A. (2002) The Mre11 complex is required for repair of hairpin-capped double-strand breaks and prevention of chromosome rearrangements. Cell, 108, 183–193.

58. UniProt, C. (2025) UniProt: the Universal Protein Knowledgebase in 2025. Nucleic Acids Res, 53, D609–D617.

59. Madeira, F., Madhusoodanan, N., Lee, J., Eusebi, A., Niewielska, A., Tivey, A.R.N., Lopez, R. and Butcher, S. (2024) The EMBL-EBI Job Dispatcher sequence analysis tools framework in 2024. Nucleic Acids Res, 52, W521–W525.

60. Robert, X., Guillon, C. and Gouet, P. (2025) FoldScript: a web server for the efficient analysis of AI-generated 3D protein models. Nucleic Acids Res, 53, W277–W282.

61. Meng, E.C., Goddard, T.D., Pettersen, E.F., Couch, G.S., Pearson, Z.J., Morris, J.H. and Ferrin, T.E. (2023) UCSF ChimeraX: Tools for structure building and analysis. Protein Sci, 32, e4792.

