## Supplemental Figures for "Replication fork remodeling proteins, Smc5/6 and Rtt107, promote palindrome-mediated genome instability"

Supplemental Figure 1

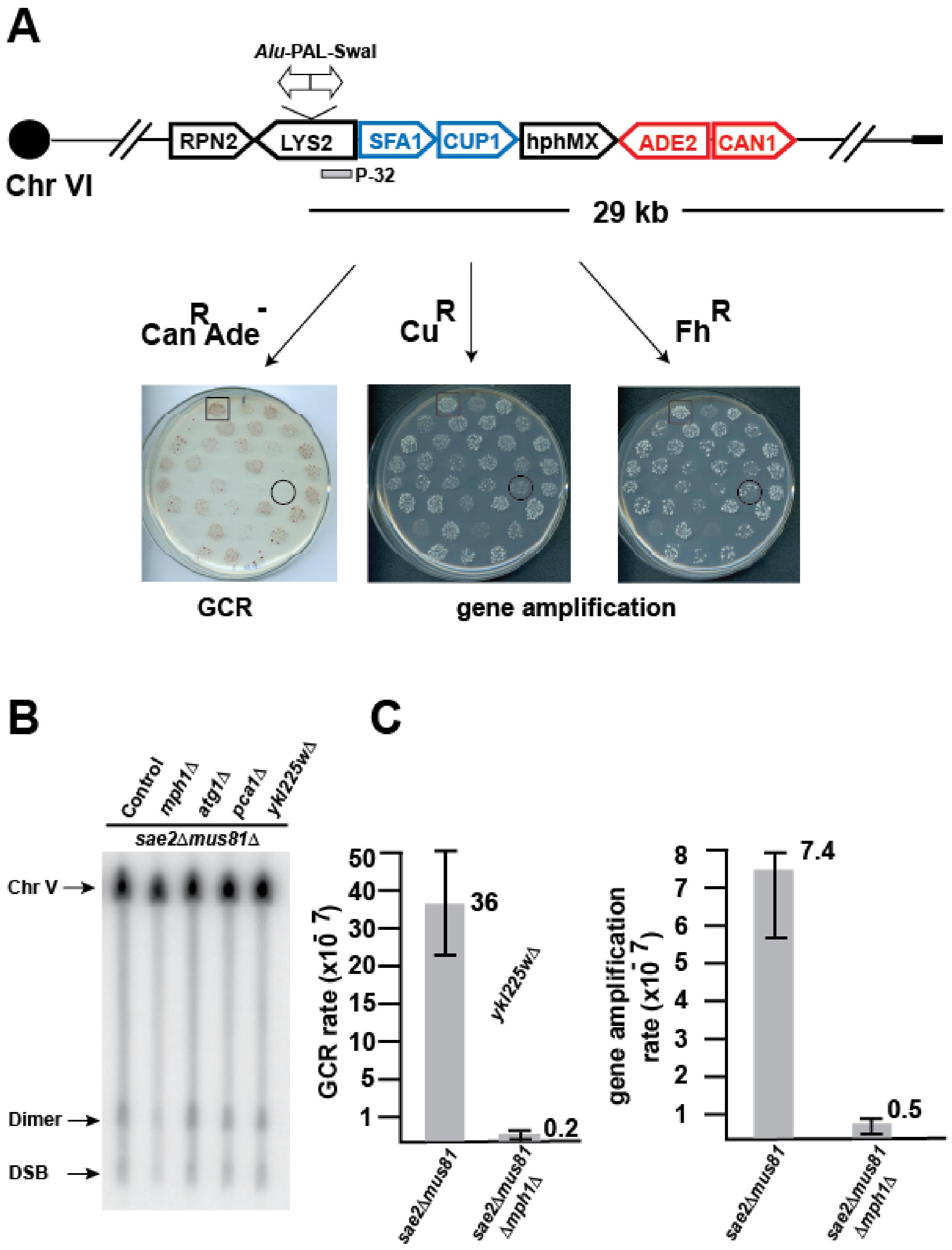

### Supplemental Figure 2

**A**

| | | Location | Structure | Conservation | Residues in other yeast species |
|---|---|---|---|---|---|
| *smc5-A6* | Q384R | Arm | α-helix | high similarity | Q or R |
| | L529P | Hinge | α-helix | low similarity | L |
| | K599E | | | | K or R |
| | H616R | | β-sheet | | H |
| | N665Y | Arm | α-helix | | N |
| | L761M | | | | Mostly L |
| | ***L897Q*** | | | high similarity | L |
| *smc5-B18* | C504R | Hinge | α-helix | low similarity | C |
| | K533R | | linker | | K |
| | V993A | Head | α-helix | high similarity | V |
| | ***C20Δ*** | | N. A. | low similarity | N. A. |
| *smc6-S9* | I830V | Arm | α-helix | high similarity | I |
| | E848G | | | low similarity | Mostly E |
| | ***P1038L*** | Head | linker | low similarity | P |
| | E1105F | | | low similarity | E |

**B**

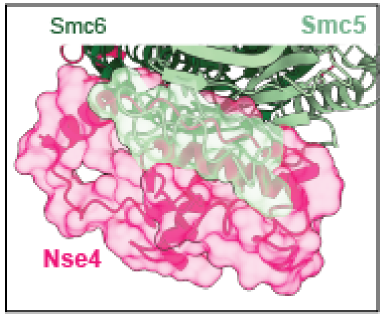


**D**

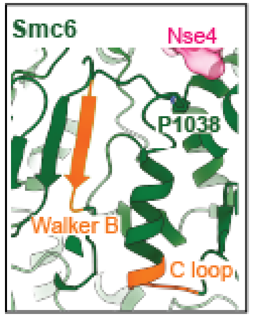


**C**

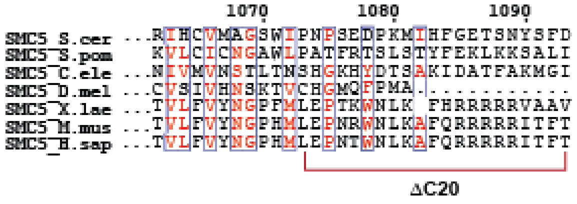


**E**

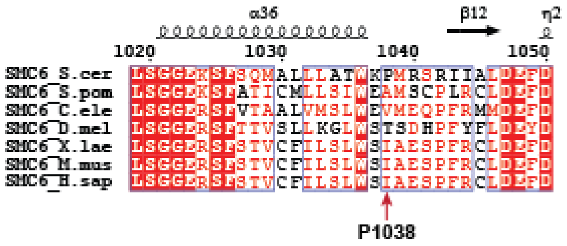
